# 22-hydroxycholesterol leads to efficient dopaminergic specification of human Mesenchymal Stem Cells

**DOI:** 10.1101/250050

## Abstract

In the current study, we aim to investigate the neurogenic effect of 22 (R)- hydroxycholesterol (22-HC), on hMSCs obtained from bone marrow, adipose tissue and dental pulp. The effect was evaluated on the basis of detailed morphological and morphometric analysis, expression of genes and proteins associated with maturation of neurons (NF, MAP2, TUJ1, TH), channel ion proteins (Kv4.2 & SCN5A), chemical functionality (DAT), synapse forming tendency of neurons (Synaptophysin) and transcription factors (ngn2 & Pitx3), efficiency of generation of dopaminergic neurons and functional assessment. The percentage of non- DA cells generation (Ach, TPH2, S100 and GFAP positive cells) was also evaluated to confirm selective neurogenic potential of 22-HC. Post analysis, it was observed that 22-HC yields higher percentage of functional DA neurons and has differential effect on various tissue- specific primary human MSCs. This study, which is one of its kinds, may help in improvising the approach of cell based treatment regimes and drug testing in pharmaceutical industry for Parkinson’s disease.

## Introduction

Cholesterol is a type of lipid, which is biosynthesized in all types of animal cells and is an important part of the cell membrane. Human brain consists of as high as ∼25% of the lipids^1^. Cholesterol, along with other lipids, self- assembles and forms various structures, contributing to the varied functions of the brain^2^. The lipidome profile of the central nervous system (CNS) is associated with neuronal activity, cognitive behaviour and various neurological disorders^3^. While cholesterol is required for dopaminergic (DAergic) neuronal maturity in terms of maintaining synapses and neurotransmission, they are also required for their survival^4–5^.

Oxysterols are the oxidized derivatives of cholesterols which show their effect in various functioning through lxr-α and lxr-β. Lxr (liver X receptors) are the nuclear receptors which function through their oxysterol ligands and control various activities like cell division, ventral midbrain neurogenesis and DAergic neurogenesis^5^. The study reported by Paolla Sacchetti et al in 2009 states the precedence of various types of oxysterols like 24-HC and 25-HC in DAergic neurogenesis in mice brain. The same study reports the relevance of 22-HC in DAergic neurogenesis in human embryonic stem cell (hESC) lines, H9 and HS181. Oxysterols have also been reported to maintain the balance between neuronal and glial cells generation.

However, till date there is no report that states the effect of oxysterols in generation of DAergic neuronal cells from Mesenchymal Stem Cells (MSCs). Lately, MSCs are being considered as very promising candidates for cell replacement therapies in various degenerative diseases, with advantages of the ease of their procurement from tissues like bone marrow (BM), adipose tissue (AD), dental pulp (DP), Wharton’s jelly (WJ), etc., maintenance in the culture, minimal ethical concerns and no immunogenicity. Hence, efficient generation of functional DAergic neurons from MSCs *in vitro* is a pre- requisite to use MSCs for translational purpose.

Our aim is to investigate the effect of 22-HC on human Mesenchymal stem cells (hMSCs), which in turn has the most optimum translational implication. Considering the specific action of 22-HC in DAergic neurogenesis and very few evident/ explanatory studies using hESCs or *in vivo* mice models, we ventured to explore its contribution towards generation of DA neurons from hMSCs.

Therefore, considering the varied roles and initial studies stating the importance of 22-HC in DAergic neurogenesis and the significance of MSCs in clinical set- up, we designed this study to coax primary MSCs obtained from various tissue sources like BM, AD and DP using 22-HC in combination with FGF2 towards DAergic neurons and their characterization at morphological, ultra-structural, transcriptional and translational levels. Induced hMSCs were also assessed for their functionality by calcium ion imaging method. Apart from the analysis of DAergic neuronal cells, presence of any other cell types was also enumerated. Changes occurring in the cells at ultra structural level were also studied. This detailed comparative analysis on a single platform has given the futuristic direction of research in the field of DAergic neurogenesis and regenerative medicine.

## MATERIALS & METHODS

### Ethics Statement

The study was conducted after receiving ethical clearance from Institutional Ethics Committee (IEC) and Institutional Committee for Stem Cell Research (IC-SCR), AIIMS, New Delhi. All the methods described in this study were performed in accordance with the relevant guidelines and regulations of the Institution.

### Cell Culture Revival and expansion of Bone Marrow- Mesenchymal Stem Cells (BM-MSC), Adipose tissue derived Mesenchymal Stem Cells (AD-MSC) and Dental pulp derived Mesenchymal Stem Cells (DP-MSC)

Cryopreserved BM- MSC, AD-MSC and DP-MSC were used for the study. Informed consent was obtained from the patients or legal representatives at the time of bone marrow, adipose tissue or extracted tooth/teeth collection for previous research projects. Age of patients ranged from 24 to 40 years for both BM- MSC and AD-MSC and 20-28 years for DP-MSC. Cells of 5 healthy donors of each MSC type, which were cryopreserved at first passage, were revived in DMEM-LG medium (Gibco, USA) with 10% fetal bovine serum (FBS) (Hyclone, USA) and penicillin (100U/ml) streptomycin (100μg/ml) (Gibco, USA) (Expansion medium), expanded and characterized. Viability of the revived cells was measured by trypan blue staining (Invitrogen, Life Technologies, USA). Followed by characterization (by surface marker profiling using flow cytometry and trilineage differentiation potential), cells from 3^rd^ to 5^th^ passage were used for all the further experiments^6^.

***Neuronal Differentiation*:** For neuronal differentiation, induction medium containing Neurobasal media (Gibco, USA), B27 supplement (Gibco, USA), EGF, FGF2 (10ng/ml each) (PeproTech Asia) and 22-HC (2μM) (Sigma, USA), L-Glutamine (Gibco, USA), PenStrep (Gibco, USA) was used. The induction protocol was carried out for 14 days with media change on every 3rd day. After termination of induction period at 14 days, the cells were used for further experiments.

### Neurites’ length Analysis

This was performed as per the already established protocol of the lab. Briefly, induced hMSCs were examined for the morphological changes under inverted microscope. Images were captured at 10X magnification and analysed using SI Viewer software (Tokyo, Japan) for the number and length of neurites, length of axon and area and diameter of the cell body. Respective uninduced hMSCs were used as experimental control^6^.

### Scanning Electron Microscopy (SEM)

Samples for SEM analysis were processed as per the established protocol of the lab^7^. Briefly, hMSCs were cultured and differentiated over cover slips. These samples on coverslips were collected and fixed with Karnovsky fixative (4% paraformaldehyde and 1% glutaraldehyde in 0.1 M Phosphate Buffer (pH 7.4)) for 6-8 hours at 4°C. Dried samplesn were mounted over aluminium stubs and sputter-coated with gold prior to imaging with EVO18 scanning electron microscope (Zeiss, Oberkochen, Germany) at 5 KVA in secondary electron imaging mode.

### Transmission Electron Microscopy (TEM)

Samples for TEM analysis were processed as per the established protocol of the lab^8^. After differentiation of hMSCs into neuronal cells, medium was removed and cells were given a gentle wash using PBS (pH 7.4), followed by fixation of cells by Karnovsky’s fixative (4% paraformaldehyde and 1% glutaraldehyde in 0.1 M Phosphate Buffer (pH 7.4)) for 6-8 hours at 4°C. After fixation, cells were washed gently with PBS. Cell number was such that they form a pellet of 100μl upon centrifugation. Water from the cells was removed by treating them with a series of ascending concentrations of the dehydrating agent, ethanol. Ethanol was cleared by treating the cells with xylene. After this, samples were dehydrated in ascending grades of acetone and embedded in araldite CY212. Thin sections (70 nm) were cut with a glass knife and mounted onto nickel grids. They were contrasted with uranyl acetate and lead citrate and viewed under a transmission electron microscope (Tecnai, G 20 (FEI)).

### Transcriptional Characterization of MSC induced into neuronal cells Quantitative reverse transcription- polymerase chain reaction (qRT-PCR)

After differentiation, total RNA from all the experimental groups was extracted by phenolchloroform method as previously described6. Single strand cDNA synthesis was performed by using cDNA synthesis kit from Life Technologies (California, USA) according to the manufacturer’s protocol. Expression of Nestin, Neurofilament (NF), Microtubule associated protein (MAP2) and Tyrosine hydroxylase (TH) was studied in both induced and uninduced MSC. All these primers were obtained from Sigma (Missouri, USA) (data not shown).

qRT-PCR experiments were performed using Realplex real time PCR detection system (Eppendorf, Germany) as previously described (Singh et al., 2017). qRT-PCR was done for Nestin, NF, MAP2, β III tubulin (Tuj1), TH and transcription factors, PitX3 and Ngn2. To study the genes related to dopamine transportation, expression of DAT (dopamine transporter) gene was studied. Apart from these, genes related to various ion channels like Kv4.2 (potassium channel) and SCN5A (sodium channel) were also studied. Expression of nuclear receptors of 22-HC, i.e., LXRα and LXRβ were also studied. Primers of qRT- PCR grade were procured from Sigma (Missouri, USA).

The expression of the genes of interest was normalized to that of the housekeeping gene, glyceraldehyde-3-phosphate dehydrogenase (GAPDH). Melting curve was used to confirm the results and data were analyzed using the graph pad prism software. Details of the primers used are given in Supplementary Table 1.

### Immunocytochemistry

The assay was performed as previously described^6^. Briefly, fixed cells were incubated overnight at 4°C with primary monoclonal antibodies against Nestin (1:400), MAP2 (1:250), TH (1:200) and DAT (1:200) (Abcam, USA). After induction period, hMSCs were washed five times with PBS and incubated with fluoro isothiocyanate (FITC) and texas red (TR) conjugated secondary antibodies (1:500 dilution) for 1 hour at room temperature. Finally, after washing five tmes with PBS, cells were counter stained with 4´,6-diamidino-2-phenylindole (DAPI) to visualize the cell nuclei. Cells were washed thrice with PBS to remove excess DAPI stain. Stained cells were examined using a fluorescence microscope equipped with a digital camera (Nikon Eclipse 80i, Japan).

### Intracellular staining for Flow Cytometry

hMSCs after induction with both the induction protocols, were labeled for Nestin (1:100), MAP2 (1:200), TH (1:100), synaptophysin (1:100), DAT (1:100), GFAP (1:100), TPH2 (1:150), Ach (1:150) and S100 (1:100) as previously described^6^. Briefly, the cells were incubated with primary antibodies (dilution of 1:100) for 1 hour 20 mins at 4 °C, followed by washing and incubation with secondary antibody labelled with fluorochrome tagged secondary antibody (dilution of 1:400) for 30 mins at room temperature. The cells were then washed and suspended in PBS and acquired on BD LSR II flow cytometer (Becton Dickinson, USA) with minimum of 10,000 events for each sample and analyzed with FACs DIVA software (version 6.1.2). All the antibodies were procured from Abcam, USA.

### Immunoblotting

Immunoblotting for the expression of neuronal cell specific proteins was performed with both induced and uninduced control cells, as previously described^6^. Briefly, after preparing whole cell lysates using RIPA buffer (Sigma, USA), the protein quantification was done using Bicinchoninic Acid (BCA) Assay method. Protein extracts (30 μg) were subjected to SDS159 PAGE using 12´ Tris/HCl SDS (Sodium dodecyl sulphate) gels and transferred onto PVDF membranes (Membrane Technologies, India). After blocking the membranes with 3´ BSA, they were incubated with primary antibody against β- actin (Abcam,UK, 1:2500), MAP-2 (Abcam, UK, 1:1500) and tyrosine hydroxylase (Santa Cruz, USA, 1:1000) in 1´ BSA163 phosphate saline buffer (PBS) overnight at 4°C. Post incubation, membranes were washed thrice in PBST and incubated with the appropriate horseradish peroxidase (HRP)-conjugated secondary antibody (1/4,000) (Dako, USA) for 2 hours at room temperature. Membranes were developed with chemiluminescence detection reagent (Pierce, USA) and acquired by using Gel Imager machine (Fluor Chem E, Cell Biosciences, Australia).

### Calcium ion Imaging

Change in the concentration of calcium ions was studied by calcium ion imaging in hMSCs induced for 12 days in all study groups, as previously described^6^. Briefly, hMSCs upon induction, were stained with 10μM of Fura red AM dye. After washing thrice with HBSS, the cells were activated using 50mM KCl solution. Time lapse recording was made at 488nm and 457nm for 3 minutes. Baseline readings were obtained before adding KCl solution to the cells. The experiment was performed using Leica Confocal Microscope (Model TCS SP8). The ratio of florescence at both the wavelengths was obtained and respective graph was plotted. The experiment was performed on 3 samples each. The data was analysed using Leica LAS AF software.

**Data Interpretation and Statistical Analysis**: Means ± SD of independent experiments were analyzed by student’s t- test, one way and two way ANOVA test (as per the requirement of data analysis). P<0.05 was considered as statistically significant. Analysis of data was done by using GraphPad Prism 5.00 software (San Diego, California, USA).

## Results

### Human Mesenchymal Stem Cells (hMSCs) require higher concentration of 22-HC for differentiation into neuronal cells

A higher dose of 22-HC is required to differentiate hMSCs into dopaminergic (DAergic) neuronal cells. As per the already reported dose of 22-HC (0.5μM −1.0μM) with hESC lines by Sacchetti et al., 2009, maximum neurogenic effect was observed at a concentration of 0.5μM of 22-HC, determined by the percentage of TUJ1 and TH positive cells. We titrated the dose of 22-HC on BM-MSC from a range of 0.5μM to 3.0μM and evaluated the results by flow cytometric enumeration of MAP2 and TH positive cells. 2 μM and 3 μM concentration of 22-HC had similar DAergic neurogenic effect, as was evident from the percentage positivity of MAP2 and TH positive cells (data not shown). Hence, it was decided to continue the induction protocol for 14 days with 2μM of 22-HC, which was higher than the earlier reported dose^5^.

### 22- HC causes neuron-like morphological changes in the human Mesenchymal Stem Cells

Upon induction of human Mesenchymal Stem Cells with FGF2 and 22-HC, basic morphological changes were observed. Typical spindle shaped morphology of MSCs started changing to the appearance of distinct cell body and cytoplasmic extensions like neurites and axon. 22-HC induced neurites- lengthening or branching hMSCs of all types. Nucleus of the cell shifted towards periphery and developed cytoplasmic extensions, through axon hillock- like structures on the cells body. The terminals of the induced cells were also observed to have multiple dendritic structures (Figure 1). These preliminary features were confirmed by scanning electron microscopic (SEM) studies. SEM studies showed development of fine neurites at the terminals of the cells post- induction period. Several fields also showed the cell to cell interaction, with extended neurites like extensions. Axon-hillock like structures was also observed clearly in the differentiated neuronal cells (Figure 2A i & ii). However, uninduced hMSCs were more flattened with spindle shape and had centralized nucleus.

**Figure. 1:**
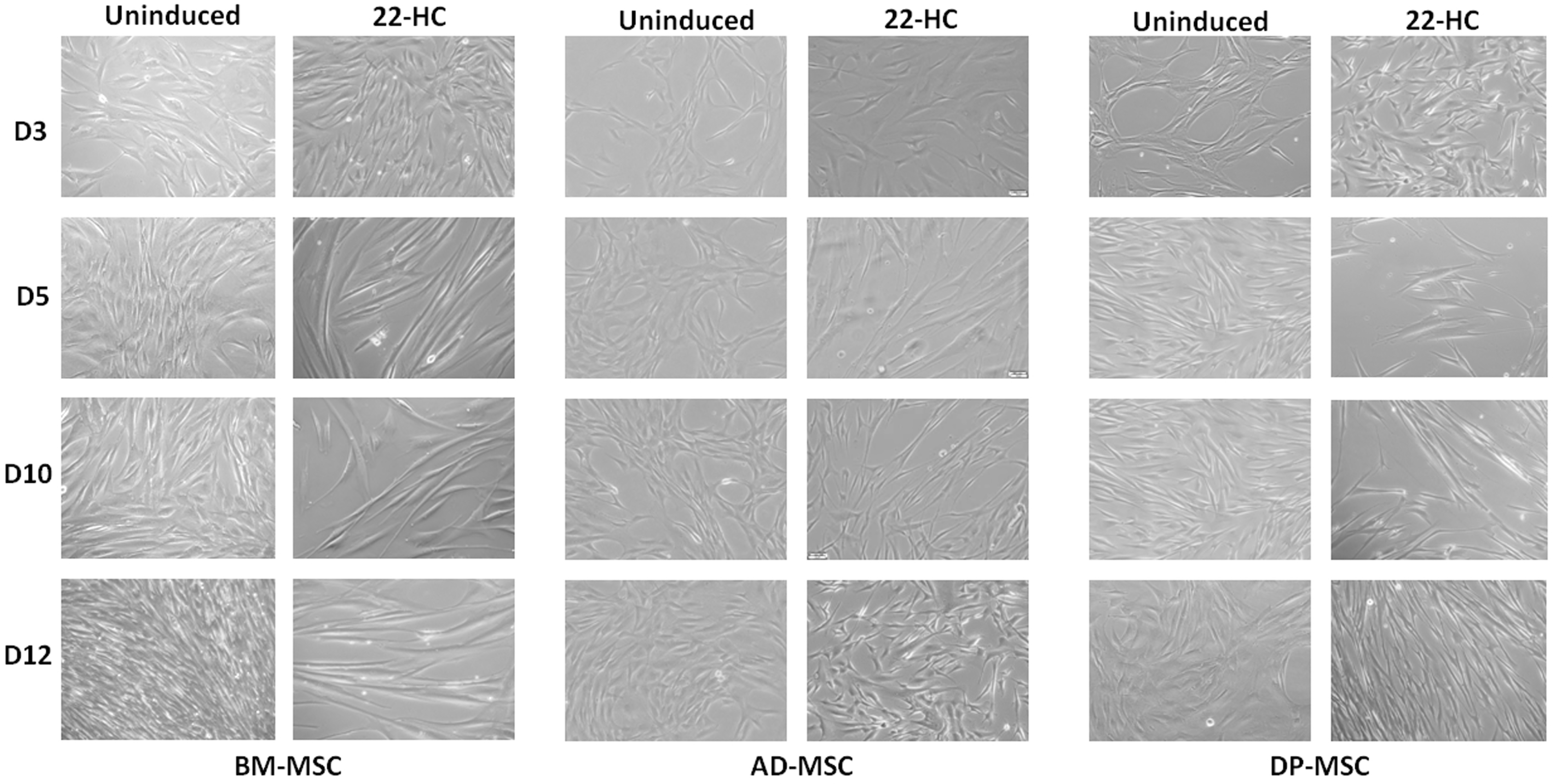
Morphological changes occurring in hMSCs during various time points (Day 3, day 5, day 10 and day 12) of neuronal induction. i) BM- MSC; ii) AD- MSC; iii) DP-MSC. The morphology of hMSCs has changed from spindle shaped to perikaryl. Appearance of neuronal morphology starts appearing from 6-7th day of induction.

**Figure. 2:**
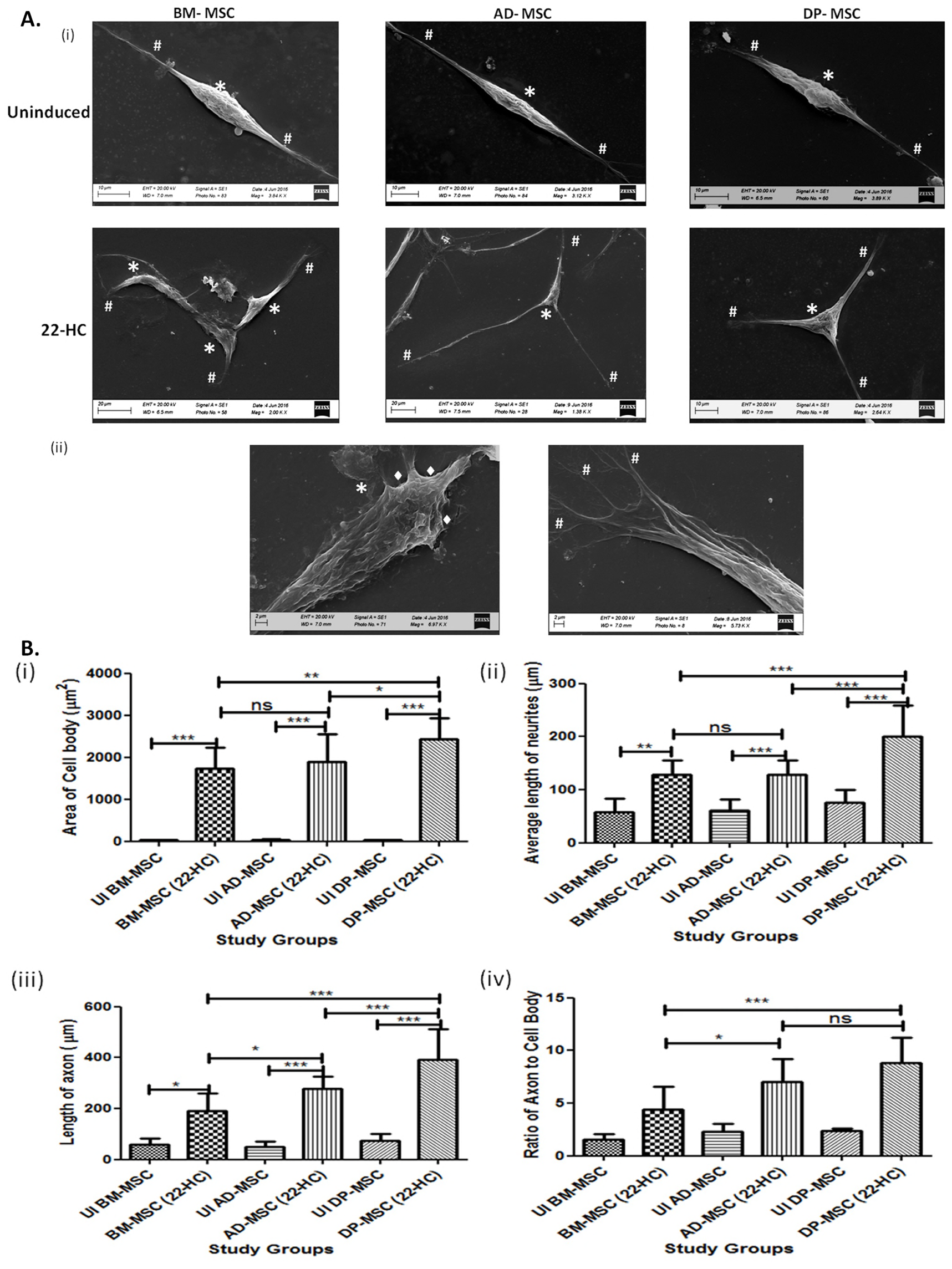
Morphological and Morphometric Characterization of differentiated hMSCs. **A. Scanning Electron Microscopic observations depicting minute morphological changes occurring in hMSCs after neuronal induction** i) Morphology of hMSCs has changed from spindle shaped to perikaryl. Terminals of the cells show appearance of minute neurites’-like structures, which enhance cell-cell interactions. Here, # denotes neuritis, * denotes cell body and •□ denotes axon hillock ii) Magnified images of differentiated cells, showing the appearance of axon- hillock, neuritogenesis and appearance of terminal neurites, facilitating cellular responses and interactions. **B. Morphometric Analysis of the neuronal cells generated in vitro by using 22-HC** i) Area of the cell body under various study groups. The graph depicts significant difference in this parameter between AD-MSC and DP-MSC; ii) Average length of neurites of differentiated cells after induction. The graph shows that the difference in the length of neurites is significantly higher in differentiated DP-MSCs, as compared to that in BM-MSCs and AD-MSCs; iii) Length of axons of cells after neuronal induction. The graph shows that the difference in the axonal length follows similar trend as average length of neurites; iv) Ratio of axon to cell body of cells under various study groups. The graph depicts that this morphological parameter of neuronal cells shows significant difference among all the three types of hMSCs under the study. For all the parameters under the study, five different samples of each type of hMSC were taken (n= 40 for each study group). Data was analysed by three independent observers.

### 22-HC induces neuronal cells- like features in hMSCs with increase in the average area of cell body, length of neurites and axon

Neuritogenesis and axonogenesis in the induced cells were assessed in detail by measuring the average area of the cell body. Average of the cell body increased from 38.51 ± 0.8036μm^2^ to 1737 ± 116.3 μm^2^ in BM-MSCs, from 43.03 ± 3.064 μm^2^ to 1897 ± 130.1 μm^2^ in AD-MSCs and from 37.85 ± 1.526 μm^2^ to 2430 ± 119.7 μm^2^ in DP-MSCs. Average length of the neurites increased from 57.59 ± 11.42 μm to 129.1 ± 6.214 μm in BM-MSCs, from 61.49 ± 6.776 μm to 128.7 ± 5.496 μm in AD-MSCs and from 75.26 ± 10.99 μm to 199.8 ± 13.56 μm in DP-MSCs. When axon length (longest neurite) was calculated, an upsurge from 57.59 ± 11.42 μm to 191.8 ± 16.03 μm in BM-MSCs, from 49.09 ± 6.913 μm to 278.5 ± 10.48 μm in AD-MSCs and from 75.26 ± 10.99 μm to 390.5 ± 25.09 μm in DP-MSCs was observed. There was no significant difference observed in the area of cell body and average length of neurites of induced BM-MSCs and AD-MSCs. On the contrary, length of axon and ratio of axon to cell body was found to be significantly higher in induced AD-MSCs as compared to those in induced BM-MSCs. However, DP-MSCs consistently had higher escalation in all the parameters studied for morphometric analysis of differentiated neuronal cells (Figure 2B i-iv).

### 22-HC leads to upregulation of DA neuronal cell traits at transcriptional level in hMSCs

Induced hMSCs were characterized for the expression of neuronal and DAergic neuronal specific genes. Post induction a relative upregulation was observed in the expression of NF (3.162 ± 0.2117, 3.274 ± 0.4632 and 29.65 ± 1.353 folds in BM-MSCs, AD-MSCs and DP-MSCs, respectively), TUJ1 (3.144 ± 0.2899, 10.91 ± 0.5304 and 18.06 ± 0.6789 folds in BM-MSCs, AD-MSCs and DP-MSCs, respectively), MAP2 (4.078 ± 0.4473, 9.072 ± 0.6825 and 35.31 ± 0.9012 folds in BM-MSCs, AD-MSCs and DP-MSCs, respectively) and TH (32.66 ± 1.775, 7.882 ± 0.8511 and 41.61 ± 0.6909 folds in BM-MSCs, AD-MSCs and DP-MSCs, respectively). However, there was no significant upregulation observed in transcriptional expression of nestin, except that in DP-MSCs with 1.344 ± 0.1671 folds (Figure 3).

**Figure. 3:**
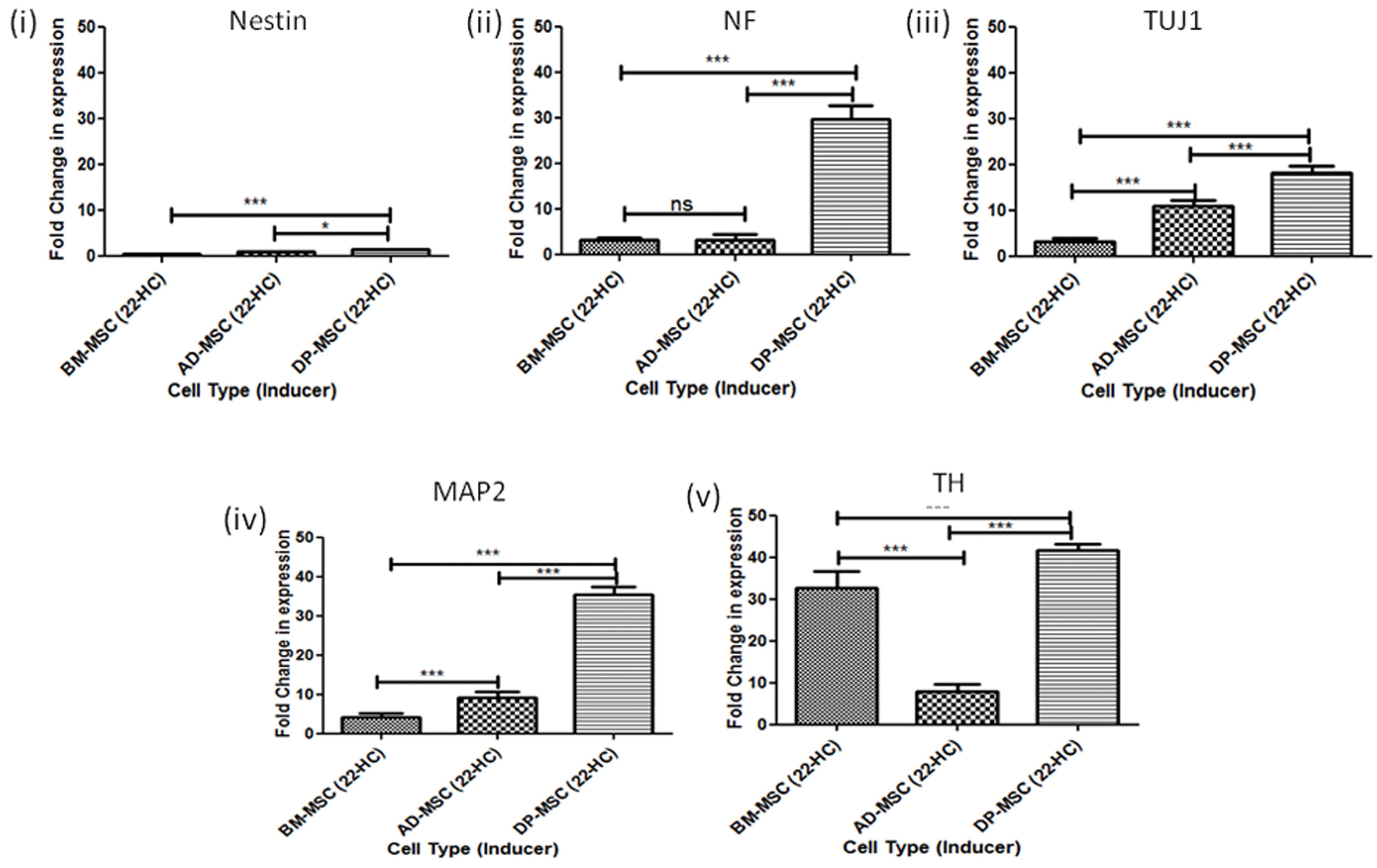
qRT- PCR transcriptional analysis of differentiated hMSCs. i) Relative fold change in mRNA expression of nestin in induced hMSCs, as compared to respective uninduced counterparts; ii) Relative fold change in mRNA expression NF in induced hMSCs, as compared to respective uninduced counterparts; iii) Relative fold change in mRNA expression of TUJ1 in induced hMSCs, as compared to respective uninduced counterparts; iv) Relative fold change in mRNA expression of MAP2 in induced hMSCs, as compared to respective uninduced counterparts; v) Relative fold change in mRNA expression of TH in induced hMSCs, as compared to respective uninduced counterparts.

Mature neuronal markers, TUJ1 and MAP2 followed similar trend, with significantly higher upregulation in AD-MSCs as compared to that in BM-MSCs. This trend was in discordance with the trend observed in rest of the genes. Hence, it may be derived that 22-HC leads to development of mature neuronal cells from AD-MSCs but does not lead to sub- specification of the same into DAergic neuronal cells, as much as it is to other hMSC types under this study.

### 22-HC causes increase in the DA neuronal cell specific proteins, corresponding to their transcriptional expression

Upon termination of induction period of 14 days, differentiated hMSCs were labelled for neuronal cell specific proteins nestin, MAP-2 and DA sub- specification protein, TH. Induced cells were found to be positive for these markers as compared to uninduced groups. There was a minimum basal expression of MAP2 & TH in uninduced hMSCs, while an upregulated expression was observed post-induction in all the hMSC types (Figure 4). An increased fluorescence intensity in the images support higher expression of MAP2 and TH in differentiated cells. However, among the various hMSCs types, highest fluorescence intensity of these protein markers was observed in case of DP-MSC. Similar trend was also observed when immunoblotting assay was performed in both uninduced and induced groups.

**Figure. 4:**
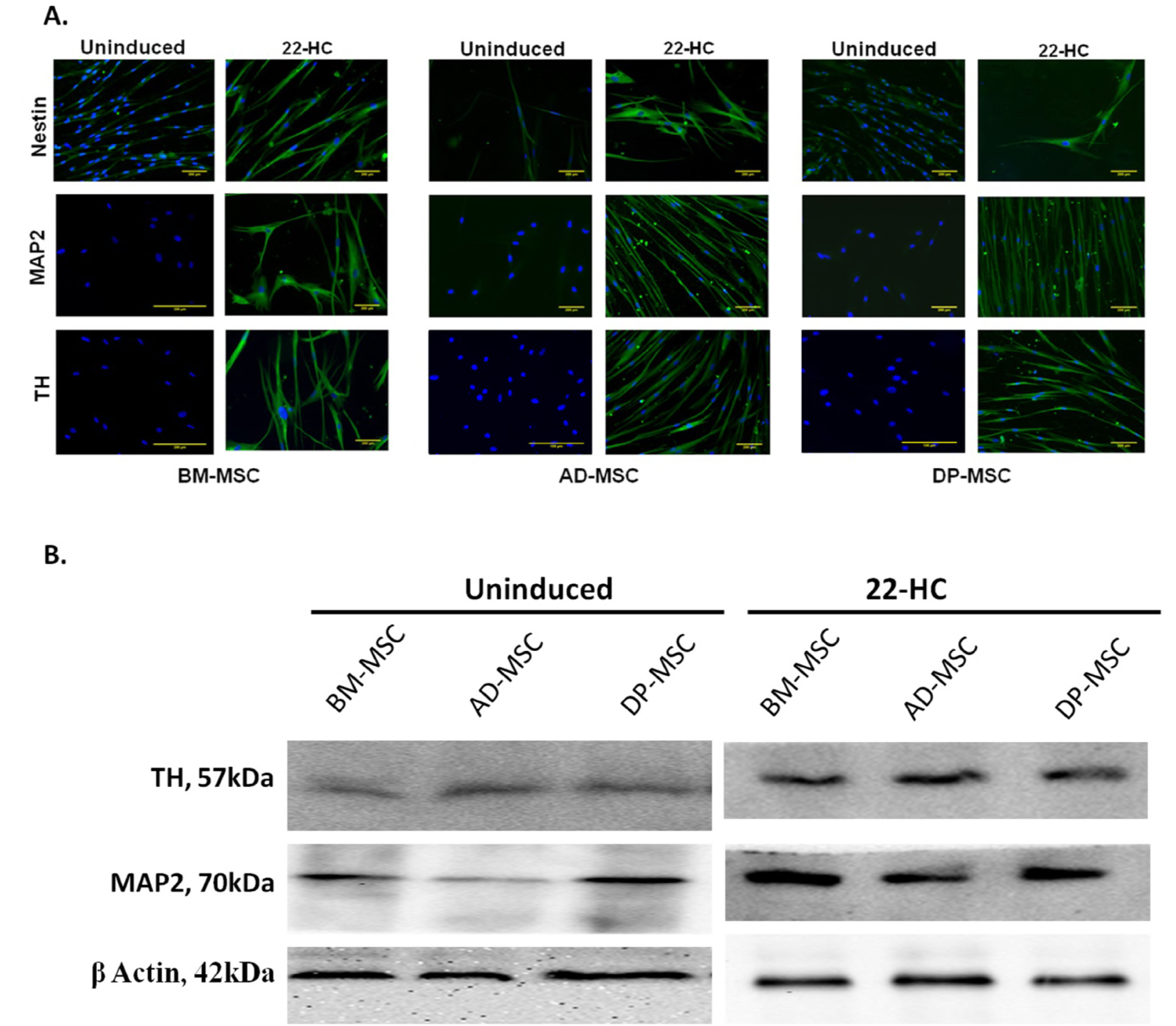
Expression of neuronal cell associated proteins in differentiated hMSCs. **A. Immunoflorescence Assay** showing expression of nestin, MAP2 and TH protein expression in hMSCs pre- and post- differentiation into DAergic neuronal cells; **B. Immunoblotting Assay** for expression of neuronal and DA neuronal cells associated proteins (MAP2 and TH) in uninduced and differentiated hMSCs.

### Induction of human Mesenchymal Stem Cells using 22-HC increases the percentage of cells positive for dopaminergic neuronal proteins

hMSCs were enumerated for the cells positive for neuronal and DAergic neuronal cells related proteins by flow cytometry. Except in the case of BM-MSCs, there was a significant increase in the percentage of cells positive for nestin (from 9.460 ± 0.964% to 15.16 ± 0.479% in AD-MSCs and from 17.84 ± 0.461 to 23.00 ± 1.145% in DP-MSCs). This observation may be attributed to the difference in the mode of action of 22-HC on the three types of hMSCs under this study, especially AD-MSCs. MAP2 positive cells were significantly increased post induction in all hMSC types, under the study. Maximum increase was observed in DP-MSCs (80.20 ± 1.544%), followed by that in BM-MSCs (67.32 ± 1.494%) and AD-MSCs (60.14 ± 0.938%). Combining the observations of transcriptional and flow cytometric analysis, it may be stated that 22-HC only led to upregulation of MAP2 gene higher in AD-MSCs as compared to BM-MSCs, but not the number of cells post- induction (Figure 5A (i & ii)).

**Figure. 5:**
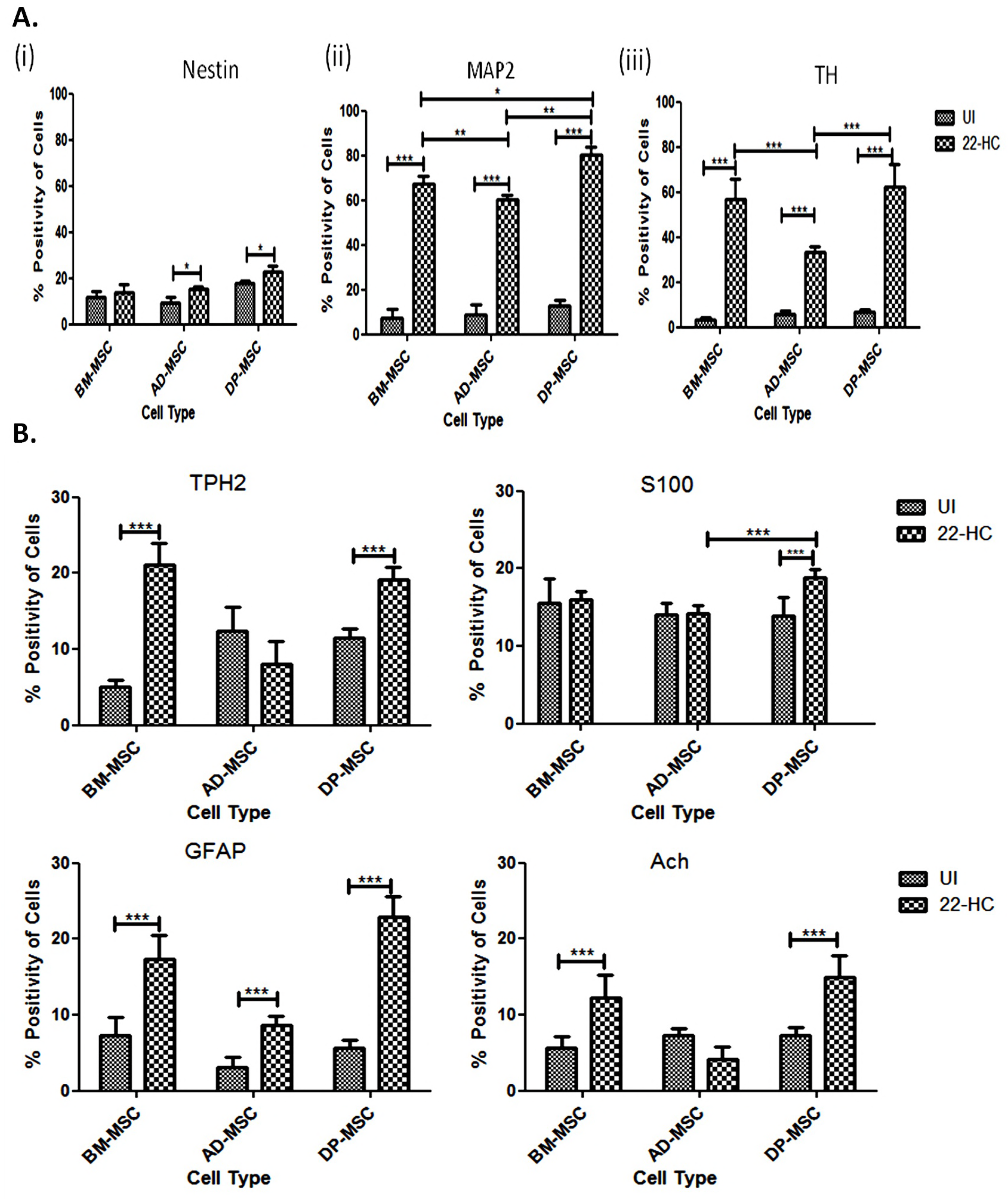
Flow cytometric analysis for cell enumeration. **A.** Graph depicting the number of nestin positive cells pre- and post- neuronal induction. DP- MSC have maximum number of nestin positive (p*, cells at the beginning of experiments, followed by those in AD-MSC and BM-MSC; ii) Graph depicting the number of MAP2 positive cells pre- and post- neuronal induction. (p*); iii) Graph depicting the number of TH positive cells pre- and post- neuronal induction. **B. Flow cytometric analysis for cell enumeration of non- DAergic neuronal cells in the cell- milieu, pre- and post- neuronal differentiation of hMSCs.**

As far as DAergic specification was concerned, TH positive cells were enumerated. No significant difference was observed in the percentage of cells positive for TH between BM-MSCs and DP-MSCs post neuronal induction (56.94 ± 4.001% and 62.32 ± 4.537%, respectively). Only 33.38 ± 1.148% cells were found to be positive for MAP2 in AD-MSCs. According to the only in vitro study by Sachhetti et al., 2009, stating the role of 22-HC in DAergic neuronal specification in hESC lines, there was an increase in TH positive cells from 22 to 60% upon treatment with 22-HC; while as high as 62.32 ± 4.537% cells were found to be positive for TH protein in DP-MSCs (considering some specific donor samples, the percentage of TH positive cells was found to be as high as 80%) (Figure 5A (iii)).

### Cell Milieu of the induced hMSCs: Presence of Non DA neuronal cells

Cell milieu of the induced hMSCs consisted of cell positive for Non- DAergic proteins like TPH2, S100, GFAP and Ach. Upon enumeration of these markers by flow cytometry, it was observed that the percentage of TPH2 increased significantly in BM-MSC and DP-MSCs only; while it decreased in AD-MSCs (however non- significantly). Similar kind of trend was observed in case of expression of Ach positive cells. However, cells positive for S100 increased only in DP-MSCs post-induction. This number was significantly higher than that in AD-MSCs.

As per the study reported by Sachhetti et al in 2009, 22-HC leads to a decrease in the GFAP positive cells when hESC lines were treated with it. However, hMSCs do not seem to follow similar trend. Instead, there was a significant increase in the cells positive for GFAP was observed in all hMSC types under study, after induction. This marker also had minimum increase in case of AD-MSCs only. These differences may be attributed to the different pathway(s), being activated by 22-HC in the various types of hMSCs under this study (Figure 5B).

### Increase in the gene expression of transcription factors responsible for maturation and survival of dopaminergic neurons

22-HC also increased the expression of transcription factors (TFs), which are responsible for maturation and survival of the DAergic neurons11. Relative fold changes in the expression level of Neurogenin 2 and PitX3 were studied in induced hMSCs, as compared to their uninduced counterparts. It was observed that while BM-MSCs and AD-MSCs showed no significant difference in the expression of PitX3, there was significantly higher upregulation of NGN2 in BM-MSCs (10.68 ± 0.6359 folds) as compared to that in AD-MSCs (8.356 ± 0.2659 folds). Whereas, DP-MSCs maintained the trend with highest upregulation of both the genes (13.75 ± 0.6226 folds of PitX3 and 15.30 ± 0.3902 folds of NGN2 in BM-MSCs, AD-MSCs and DP-MSCs, respectively). These results show that the differentiated hMSCs expressed prototypical midbrain DAergic markers at transcriptional level (Figure 6A (i & ii)).

**Figure. 6:**
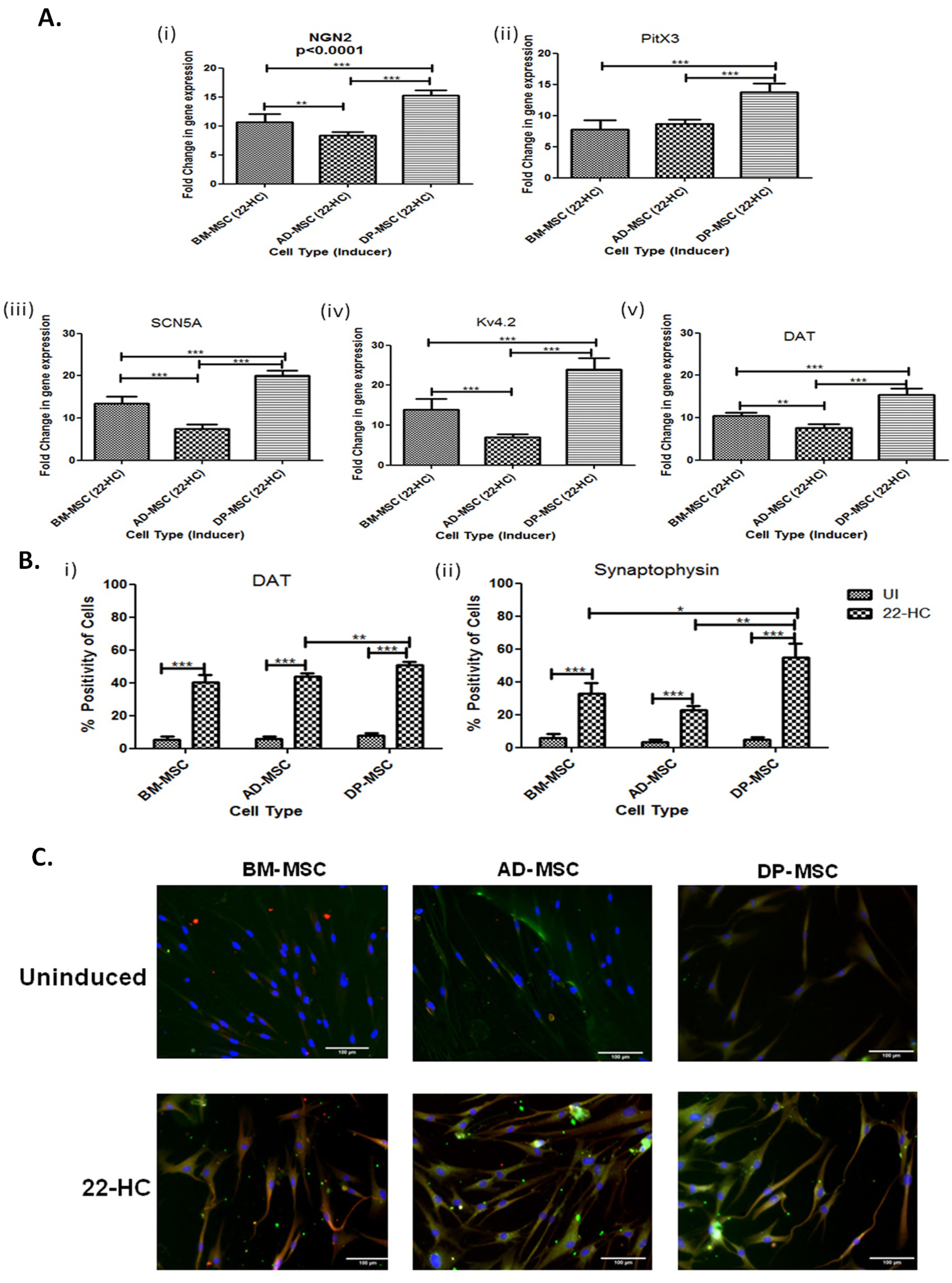
Assessment of genes and proteins associated with neuronal functionality. **A. qRT- PCR mRNA transcriptional analysis** of differentiated hMSCs for genes associated with functional behaviour of DAergic neuronal cells; **B. Flow cytometric enumeration** of number of cells positive for (i) Dopamine transporter protein, a protein responsible for release of dopamine neurotransmitter through vesicles and (ii) Synaptophysin protein which is responsible for the formation of synapse between two neuronal cells; **C. Immunoflorescence assay** to show the expression of TH and DAT proteins in hMSCs pre- and post- differentiation into DAergic neuronal cells.

### 22-HC improves functional dopaminergic specifications at both gene and protein level

Functional characterization of differentiated hMSCs was performed at gene level by studying the relative fold change in the mRNA expression level of DAT, Kv4.2 and SCN5A. A constant trend was observed where maximum upregulation of the three genes (10.47 ± 0.3574 folds DAT, 13.97 ± 1.184 folds Kv4.2 and 13.43 ± 0.7351 folds SCN5A) was highest in DP-MSCs, followed by that my BM-MSCs (10.47 ± 0.3574 folds DAT, 13.97 ± 1.184 folds Kv4.2 and 13.43 ± 0.7351 folds SCN5A) and AD-MSCs (7.542 ± 0.4570 folds DAT, 6.944 ± 0.3713 folds Kv4.2 and 7.480 ± 0.4262 folds SCN5A). Extent of upregulation of these functionality related genes was observed to be significantly lower in differentiated AD-MSCs, as compared to that in other hMSC types included in the study (Figure 6B (iii-v)).

Cells positive for DAT and synaptophysin were enumerated by flow cytometry. It was observed that as compared to their control counterparts, the number of cells in differentiated hMSCs positive for both the proteins was significantly higher. While no significant difference was observed between the outcome of BM-MSCs (40.22 ± 2.002%) and AD-MSCs (43.84 ± 0.9368%) for DAT positive cells, DP-MSCs showed significantly higher percentage of DAT positive cells (50.80 ± 0.9121%). Likewise, synaptophysin positive cells were found to be highest in number in differentiated DP-MSCs (54.98 ± 3.751%), followed by that in BM-MSCs (33.02 ± 2.713 %) and AD-MSCs (22.82 ± 1.063%). The difference was significant among all the induced cell types (Figure 6B (i & ii)).

Likewise, immunoflorescence staining revealed the presence of both TH and DAT proteins in all the hMSC types post induction. However, no or very little expression of TH protein could be observed in respective hMSC counterparts (Figure 6C).

### Ultra- structural changes, contributing towards functional DA neuronal cell specification were observed

Neuronal differentiation of hMSCs is associated with the ultra- structural modifications in the mitochondria^12^, dense core vesicles or granules (DCVs)^13^, rough endoplasmic reticulum (RER) (Wu et al., 2017), cytoplasmic filamentous condensation^14–15^and endocytotic vesicles^16^.

Mitochondria are necessary for neural development, survival, activity, connectivity, neuronal movement, axonal pruning and axonal elongation. Several neuro-developmental and neuro-degenerative diseases are associated with mitochondrial dysfunction. Mitochondrial biogenesis was increased in the *in vitro* differentiated DAergic neuronal cells, as is evident by the increase in the number of mitochondria, with globular cup-like structure and evident cristae.

There was an increase in the number of DCVs (specialized secretory organelles which are rich in ADP, ATP, ionized calcium, histamine and neurotransmitters) and RER (extend for several microns as linking network throughout highly branched dendrites and help in axonogenesis and dendritogenesis), attributing towards the increased functionality of the neuronal cells. The increase in the number of DCVs and RER is representative of the increased functionality of the cells. More DCVs may be associated with the chemical functionality of the DAergic neurons in terms of secretion of various neurotransmitters and RER may be associated with the formation of proteins required for enhanced functioning of the cell.

Cytoskeletal rearrangement controls cell morphology, regulates growth cone motility (contracted area of the neuronal cell from where neurites originate) and axon guidance and forms scaffold for transport of mitochondria and other organelles. Cytoskeletal condensation is an indicative of neuritogenesis and axonogenesis. Cytoskeletal condensation was observed in all types of hMSCs post differentiation. Microtubules were observed to be arranged in a synchronised pattern after differentiation of hMSCs with 22-HC.

Apart from these observations, small endocytotic vesicles were also observed in the ultra-structural study of hMSCs post- induction. These vesicles are an indicative of increased neuronal functionality, as they help in neuronal signalling to regulate cell survival, axon growth and guidance, dendritic branching and cell migration (Figure 7A).

**Figure. 7:**
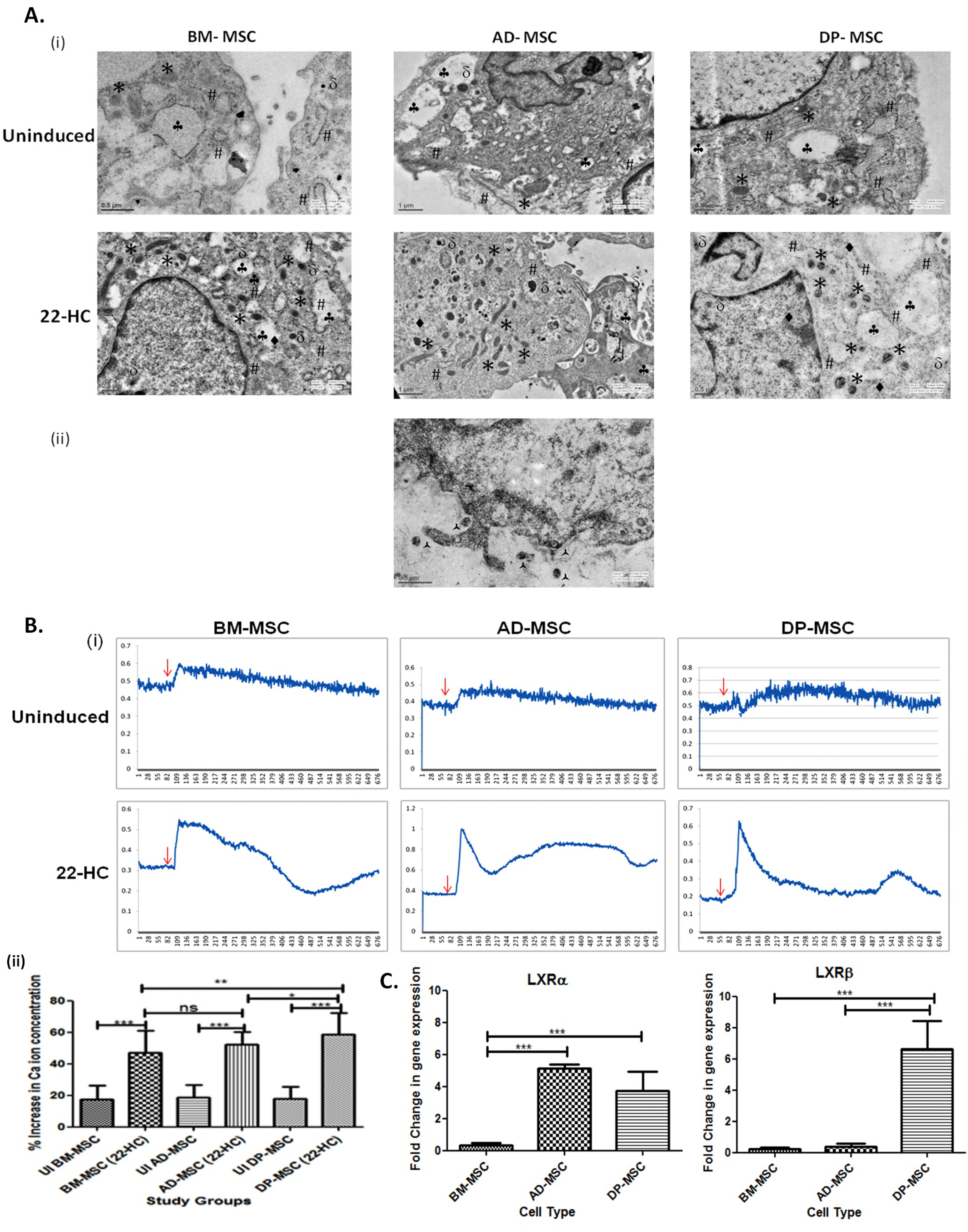
**A. Transmission Electron Microscopic observations depicting ultra- structural changes occurring in hMSCs after neuronal induction** i) Ultra- structural composition of hMSCs has changed on various parameters. There was observed increased mitogenesis, increase in dense core vesicles, rough endoplasmic reticulum, cytoskeletal condensation and endocytotic vesicles. The genesis of these cellular organelles may be associated with the increased functionality of the terminally differentiated hMSCs. ii) Part of the cell membrane showing the release of vesicles (probably synaptic vesicles) by exocytosis, contributing to the chemical functionality of the DAergic neuronal cells. In this figure, # represent rough endoplasmic reticulum, * represent mitochondria, □ represent endocytotic vacuoles, • represent cytoskeletal condensation and □ represent dense core vesicles. å represent the exocytotic vesicles released from differentiated DP-MSCs. **B. Calcium ion imaging analysis by Fura red- AM ratiometric dye:** i) Line graphs showing the changes in the Ca2+ transients upon depolarization with KCl in cells of all study groups. The uninduced hMSC did not show any change in the intracellular Ca2+ concentration after depolarization. The red arrows indicate the time point of addition of KCl for depolarization in the cell culture; ii) Graph showing the change in the percentage increase in the calcium ion concentration in hMSCs obtained from various tissue sources after DAergic neuronal induction by 22-HC. **C. qRT-PCR transcriptional Analysis of liver X receptors α and β**, which are the main nuclear receptors, reported till date, responsible for DAergic neurogenesis by oxysterols (22-HC). Our results suggest a prominent role of LXRβ in DAergic neurogenesis with 22-HC as inducer DP-MSC only. LXRα does not seem to play much distinct role in this process. However, least upregulation of both of these genes was observed in BM-MSCs.

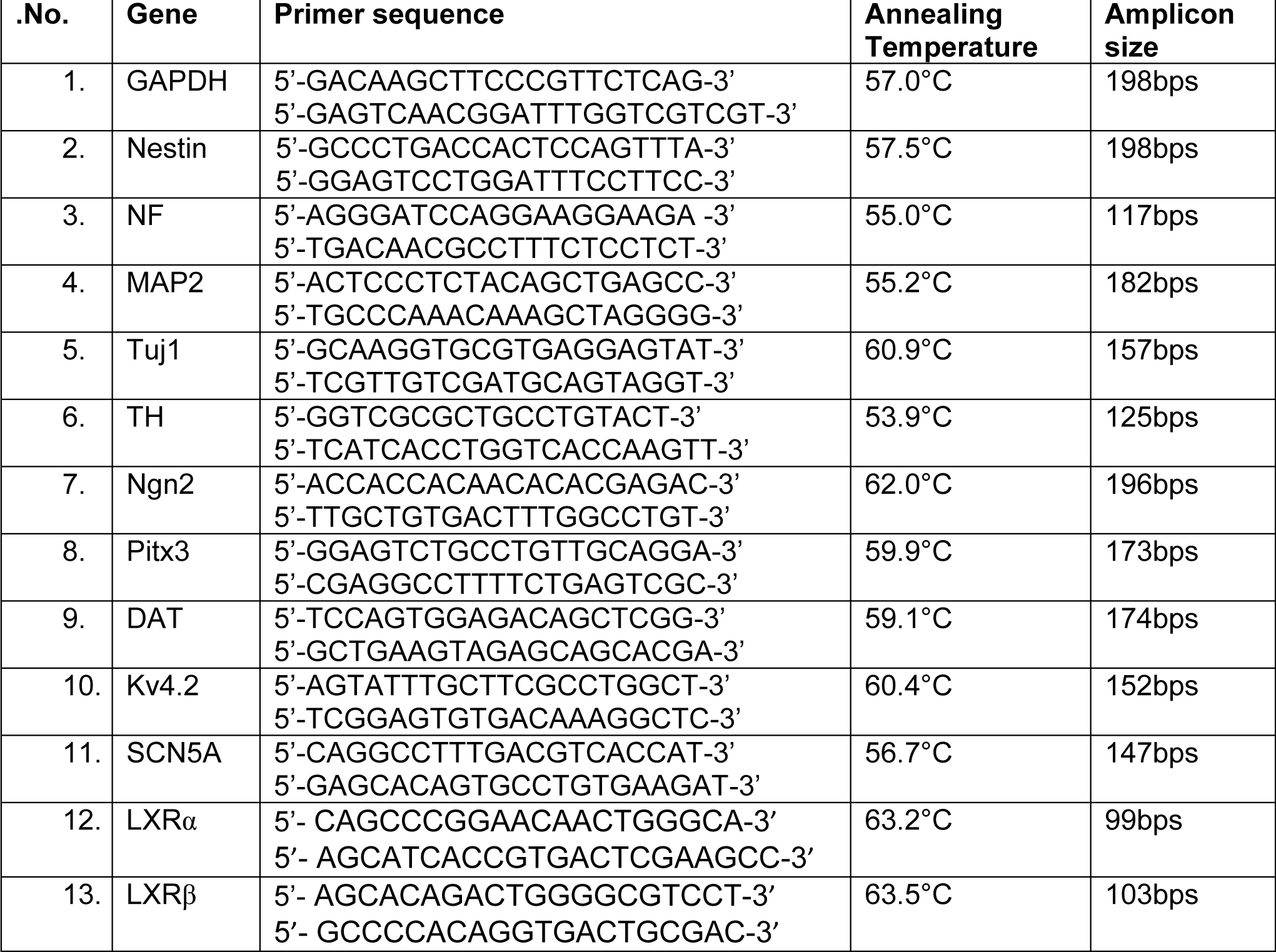

### Human Mesenchymal Stem Cells show increase in the calcium ion efflux upon treatment with 22-HC

Upon treatment with 50mM of KCl, change in the calcium ion concentration in the cytosol of the *in vitro* generated neuronal cells was observed. There was significant difference in the intracellular calcium ion transients in DAergic neuronal cells generated by treatment with 22- HC in all types of hMSCs, as compared to that in respective uninduced MSCs. In DP-MSCs the change in calcium ion transients was observed to be maximum (58.53 ± 2.622%) as compared to the control (17.93 ± 1.726%). This was followed by that in AD-MSCs (52.07 ± 1.831% in differentiated and 18.75 ± 1.711% in control) and least calcium ion transients were observed in BM-MSCs (46.98 ± 2.566% in differentiated and 17.57 ± 1.912% in control). The change in the calcium ion concentration in various study groups have been detailed in the Figure 7B (also see supplementary materials for respective videos).

### Liver X receptors (α and β) acknowledge 22-HC differently with different types of hMSCs

LXRα and LXRβ are the nuclear receptors, activated by oxygenated derivatives of cholesterol (22-HC being the natural ligand of LXRs), to accomplish various cellular functions like homeostasis of fatty acid, cholesterols and glucose metabolism. Apart from this, some studies support their role in ventral midbrain dopaminergic neurogenesis. These facts led us to investigate the fold change in the transcriptional expression of both LXRα and LXRβ in hMSCs after treatment with 22-HC for DAergic neuronal differentiation. It was observed that these receptors had different level of expression in different hMSC types after differentiation. AD-MSCs showed highest upregulation of LXRα (5.113 ± 0.1272 folds), followed by that in DP-MSCs (3.710 ± 0.6238 folds), with almost negligible changes in case of BM-MSCs. On the contrary, LXRβ was maximally expressed in DP-MSCs (6.645 ± 0.9002 folds), followed by that in AD-MSCs and BM-MSCs, with negligible upregulation (Figure 7C).

This difference in the expression of both the receptors in different hMSC types may be attributed to their origin, chief function(s) in the living system and pathway(s) that might have been activated, resulting in DAergic neurogenesis.

## Discussion

Evidences demonstrating the role of oxysterols and their nuclear receptors, LXRα and LXRβ in human midbrain dopaminergic neurogenesis have been reported by Sacchetti et. al., 2009. This study was performed on mouse embryo and human embryonic stem cell lines. It had been observed that after adding 22-HC in the induction cocktail, the percentage of cells positive for TH was increased from 22% to 60% with the subsequent decrease in GFAP positive cells. Thus, the study reports the usage of 22-HC for differentiating hESCs *in vitro* and prospects their use in regenerative medicine and drug testing in future.

However, use of hESCs in clinical set up is not recommended at present by several researchers and clinicians due to their immunogenicity and tendency to form teratoma. Hence, MSCs are preferred over hESCs for both clinical and drug testing purposes. Considering the translational aspect of the type of stem cells, in our current study, we have, for the very first time, investigated the effect of 22-HC on DAergic neuronal differentiation of human MSCs, derived from human bone marrow, adipose tissue and dental pulp. The investigation included detailed morphological, morphometric, transcriptional, translational, ultra structural and functional characterization of DAergic neuronal cells derived from these various types of hMSCs under the study.

We, hereby, report the use of 22-HC as a novel inducer for efficient *in vitro* generation of DAergic neuronal cells from human MSCs, and their detailed multi- factorial characterization. Our protocol reports as high as 80% MAP2 positive cells after induction of DP-MSCs with 22-HC. This is the highest percentage of *in vitro* generated mature neurons reported till now, using any of the established protocol^17–24^. Also our protocol yields the maximum number of TH positive DAergic neuronal cells (72% in case of DP-MSCs), as compared to any of the studies reported till date^5,6,9,14,15^. These statistics are the highest among all the reported studies till date, to the best of our knowledge. DP-MSCs showed maximal efficiency of *in vitro* generation of DAergic neuronal cells, followed by BM-MSCs and least being in AD-MSCs. The protocol presented in the current study is not only the most efficient, but also cost- effective, as it includes use of only two inducers for generation of DAergic neuronal cells *in vitro* from hMSCs that can be used for translational purpose.

We, for the very first time, report the ultra structural changes occurring in hMSCs upon *in vitro* differentiation into DAergic neuronal cells. While we have commented on the fine structural changes that appear upon differentiation of hMSCs into neuronal cells, we have also studied in detail the metamorphosis occurring in the cellular components after differentiation.

The study also hints that LXRβ plays prominent role in DAergic neurogenesis of hMSCs as compared to LXRα. However, further detailed investigations are required to prove this hypothesis. Also, the noteworthy observations in the difference of transcriptional expression of LXRα in case of AD-MSCs may be reasoned with their constitutive function of fat storage and synthesis of estrogens and androgens^24^ in the living system. We hypothesize that LXRα is more upregulated in AD-MSCs, to activate their default pathway(s). As no direct or indirect proof is available to state the exact role of these two receptors in DAergic neurogenesis and differential effect on various types of embryonic or adult stem cells, our hypothesis, needs further experimental validation.

In conclusion, our study provides comprehensive and robust evidences to state the role 22-HC in generating functional DAergic neuronal cells from hMSCs of varied origins. These in vitro generated cells showed DAergic neuronal specifications at morphological, morphometric, ultrastructural, transcriptional, translational and functional levels, with tremendous clinical and pharmacological application. As reported by Sacchetti et al in 2009, functions of LXRs are conserved in human cells. Our study gives the first evidence that oxysterols (22-HC) causes DAergic neurogenesis and leads to their morphological, genetic and functional changes. Our report has also paved the path to investigate the disparate effect of 22-HC on the three types of hMSCs taken under the study. AD-MSCs have shown off beat expression of LXRα as compared to that of LXRβ. These pathways need to be investigated further to have an insight of the mode and mechanism of action of 22-HC for midbrain DAergic neurogenesis.

## Acknowledgements

The authors would like to thank Dr. Shantanu Chowdhary and Mr. Manish Kumar, Institute of Genomics and Integrative Biology, New Delhi, India, for providing guidance and assistance in performing confocal live cell imaging (calcium ion imaging) experiments, included in this manuscript. This research work, being a part of Ph.D. thesis, has been funded by the departmental grants under Department of Cardiothoracic and Vascular Surgery, AIIMS, India. First author (Ph.D. student) was supported by University Grants Commission (UGC), Government of India during this research.

## Authors’ Contributions

Basic stem cell characterization studies: MS. Conceived, planned and performed the experiments: MS and SM; Electron microscopic studies (conceptualization, performance and analysis): SM, AKD and MS; Data Analysis: MS and SM; Contributed reagents/materials/analysis tool: SM and MS; Arrangement of research funds: BA and SM; Functional studies: MS and SM; Manuscript writing: MS and SM.

## Additional Information

Not applicable

